# The development of aperiodic and periodic resting-state power between early childhood and adulthood: new insights from optically pumped magnetometers

**DOI:** 10.1101/2024.05.13.593983

**Authors:** Marlee M. Vandewouw, Julie Sato, Kristina Safar, Natalie Rhodes, Margot J. Taylor

**Affiliations:** Department of Diagnostic Imaging, Hospital for Sick Children, Toronto, Canada; Program in Neurosciences & Mental Health, Hospital for Sick Children, Toronto, Canada; Autism Research Centre, Bloorview Research Institute, Holland Bloorview Kids Rehabilitation Hospital, Toronto, Canada; Institute of Biomedical Engineering, University of Toronto, Toronto, Canada; Sir Peter Mansfield Imaging Centre, School of Physics and Astronomy, University of Nottingham, Nottingham, United Kingdom; Department of Medical Imaging, University of Toronto, Toronto, Canada; Department of Psychology, University of Toronto, Toronto, Canada

**Keywords:** optically pumped magnetometers, magnetoencephalography, development, power, aperiodic activity, periodic activity

## Abstract

Neurophysiological signals, comprised of both periodic (e.g., oscillatory) and aperiodic (e.g., non-oscillatory) activity, undergo complex developmental changes between childhood and adulthood. With much of the existing literature primarily focused on the periodic features of brain function, our understanding of aperiodic signals is still in its infancy. Here, we are the first to examine age-related changes in periodic (peak frequency and power) and aperiodic (slope and offset) activity using optically pumped magnetometers (OPMs), a new, wearable magnetoencephalography (MEG) technology that is particularly well-suited for studying development. We examined age-related changes in these spectral features in a sample (*N*=65) of toddlers (1-3 years), children (4-5 years), young adults (20-26 years), and adults (27-38 years). Consistent with the extant literature, we found significant age-related decreases in the aperiodic slope and offset, and changes in peak frequency and power that were frequency-specific; we are the first to show that the effect sizes of these changes also varied across brain regions. This work not only adds to the growing body of work highlighting the advantages of using OPMs, especially for studying development, but also contributes novel information regarding the variation of neurophysiological changes with age across the brain.

## 1 Introduction

The study of the development of neurophysiological brain signals in health and disease has a long and rich history. Neural oscillations, or the periodic fluctuations in neuronal activity, have been the primary focus, with a wealth of investigations into their role in sensory and cognitive processes, and on their marked evolution across the lifespan. While most studies have used electroencephalography (EEG), there has been increasing interest in magnetoencephalography (MEG) due to its comparable temporal and superior spatial sensitivity (Baillet, 2017; Hämäläinen & Lundqvist, 2019). However, both EEG and traditional MEG systems pose several challenges for studying the development of neural oscillations (R. M. Hill et al., 2019), particularly when examining both toddlers and adults. Periodic activity is also only one aspect of neurophysiological signals, with an increasing body of work suggesting that the background activity in the brain, or the aperiodic activity, also undergoes significant developmental changes (Voytek et al., 2015) and has distinct cognitive relevance (Thuwal et al., 2021). Here, we are the first to use optically pumped magnetometers (OPMs), a new, wearable MEG technology that is more suitable for studying very young children, to characterize the changes in both periodic and aperiodic neural activity between toddlerhood and adulthood.

Periodic activity is defined by narrow peaks of power in the frequency domain, ranging from slow frequencies such as delta through to high gamma (Buzsáki & Draguhn, 2004). In contrast, aperiodic activity does not contain rhythmic oscillations, instead, it is “scale-free”, meaning it does not contain a dominant temporal scale (B. J. He, 2014). Its distribution follows the “1/f” power law – exponentially decreasing power with increasing frequency – which can be characterised by its slope and offset (Gao et al., 2017). While many studies have attempted to link periodic or oscillatory activity with a wide range of disease states over the years, reliable biomarkers have remained elusive (e.g., (Parellada et al., 2023)). It is suggested that the aperiodic component may be more stable and sensitive to a range of clinical conditions (Pani et al., 2022).

Importantly, aperiodic activity has been proposed as a means of determining the excitatory-inhibitory (E/I) balance in the brain (Gao et al., 2017) that has been implicated in a range of developmental conditions (see (Pani et al., 2022) for a review). However, before it can be used clinically, a better understanding of aperiodic activity across development is required.

Investigations into the development of aperiodic activity are in their infancy and have predominantly relied on EEG, with limited work using MEG (e.g., (Manyukhina et al., 2022)). These studies have shown that the aperiodic slope and offset both decrease with age within infancy (Schaworonkow & Voytek, 2021), childhood (A. T. Hill et al., 2022; McSweeney et al., 2023), and adolescence (McSweeney et al., 2021), as well as extending across these developmental periods (Cellier et al., 2021; Favaro et al., 2023; Tröndle et al., 2022). On the other hand, while age-related changes in neural oscillations have been well characterised (e.g., (Ebersole & Pedley, 2003)), traditional analyses of neural oscillations within canonical frequency bands can be confounded by aperiodic activity (Donoghue, Dominguez, et al., 2020). For example, frequently reported decreases in alpha power in aging were found to be absent when controlling for the aperiodic slope (Cesnaite et al., 2023; Merkin et al., 2023). Thus, it is important to re-examine developmental changes in periodic activity while accounting for the influence of its aperiodic counterpart. A small selection of studies has done so, finding age-related increases in aperiodic-adjusted peak frequency in alpha (Cellier et al., 2021; McSweeney et al., 2023; Tröndle et al., 2022), but not beta (A. T. Hill et al., 2022). Findings were mixed with respect to aperiodic-adjusted alpha power (Cellier et al., 2021; A. T. Hill et al., 2022; Tröndle et al., 2022), however, a very well-powered sample reported increases across childhood into early adulthood (Tröndle et al., 2022).

A confound with these EEG studies, however, is that brain maturation occurs synchronously with skull thickening, which would lead to greater resistance and hence reduced EEG amplitudes (Tröndle et al., 2022). Therefore, changes in the aperiodic component seen with age could be in part secondary to physical skull changes irrelevant to brain function. The skull and scalp also limit the spatial resolution of EEG to approximately 1 cm, and thus developmental changes have been restricted to examinations across the whole brain or across lobes. On the other hand, MEG signals pass through the skull and scalp with little distortion, removing that confound while also allowing for far better spatial resolution of the generating sources of activity, to the order of a few millimetres (Hari & Salmelin, 2012). OPM-MEG specifically offers a significant advantage for studying development compared to traditional MEG systems using superconducting quantum interference devices (SQUIDs; (R. M. Hill et al., 2019)).

Compared to SQUID-MEG, OPM-MEG is more tolerable to head motion, a well-established age-related confound. SQUID-MEG helmets are also “one-size-fits-all”, which means there are coverage, signal strength, and signal-to-noise ratio differences between smaller and larger heads, which is a significant concern for developmental studies. On the other hand, in OPM-MEG the sensors can be mounted on a helmet that is customized to an individual’s head size which can mitigate these issues (Rier et al., 2024).

In this study, we are the first to investigate the development of aperiodic and periodic neural signals using OPM-MEG. We examined age-related changes in the aperiodic slope and offset alongside adjusted peak frequencies and power using a sample of very young children (1 – 5 years of age) and adults (20 – 38 years of age). Leveraging the spatial resolution of MEG, we characterized the developmental changes within individual brain regions, alongside whole-brain analyses.

## 2 Materials and methods

### 2.1 Participants

Seventy-three participants (20 toddlers (1 – 3 years), 16 young children (4 – 5 years), 18 young adults (20 – 26 years), and 19 adults (27 – 38 years)) were recruited as part of a larger study at the Hospital for Sick Children. All participants were typically developing without a current diagnosis or history of a neurological or neurodevelopmental disorder, chromosomal or major congenital abnormality. Individuals were eligible for the current study upon successful completion of the resting-state *Inscapes* paradigm (described in the following section). Written informed consent was provided by the caregiver or participant for the children and adults, respectively, and the study protocol was approved by the Hospital for Sick Children research ethics board.

### 2.2 OPM acquisition

A 40 dual-axis (80 channels) OPM system ((R. M. Hill et al., 2022); QuSpin Incorporated, Colorado, USA; Cerca Magnetics Limited) with a 1,200 Hz sampling rate was used to collect data while participants watched 5-minutes of the *Inscapes* resting-state paradigm (Vanderwal et al., 2015), which consists of a movie of slowly moving shapes and accompanying piano music. Before presenting the paradigm, a 5-minute empty-room noise recording was also obtained. OPM sensors were mounted in one of four possible 3D-printed helmets of varying sizes (Cerca Magnetics Ltd.; three of the four helmets were used in the current study). For each participant, the helmet choice was customized to their head circumference. The participant wore the helmet while seated within a magnetically shielded room (Vacuumschmelze, Hanau, Germany). Bi-planar coil panels and OPM reference sensors (QuSpin Incorporated) were positioned on each side of the participant for dynamic and static nulling of the background magnetic field and its drift, maintaining sensor operation between ±3.5nT (Holmes et al., 2019; Rea et al., 2021). A four-camera system (OptiTrack Flex 13, NaturalPoint Incorporated, Oregon, USA) with infra-red markers placed on the helmet was used to continuously track head movement; head motion data was not successfully acquired for three participants. For co-registration of the OPM data with brain anatomy (R. M. Hill et al., 2020; Zetter et al., 2019), digitisations of the participant’s head with the helmet were acquired using a 3D optical imaging system (Einscan H, SHINING 3D, Hangzhou, China).

### 2.3 Preprocessing

All preprocessing was performed using an OPM preprocessing pipeline developed in-house using the FieldTrip toolbox (version 2202-02-14; (Oostenveld et al., 2011) implemented in MATLAB (version R2021a; (The Mathworks Inc., 2018)). For both the resting-state and empty-room noise recordings, noisy channels were removed using an outlier detection algorithm (Safar et al., 2024), and homogeneous field correction was used to suppress interference from sources outside the head (e.g., environmental noise; (R. M. Hill et al., 2022; Tierney et al., 2021)). After bandpass filtering (1-150Hz, 4th order, two-pass Butterworth), peaks present in the power spectra of both the empty-room and resting-state data were identified as noise and were subsequently band-stop filtered (4th or 3rd order, two-pass Butterworth). The resting-state data were epoched into 1-second segments, and epochs with signals exceeding an artefact rejection threshold were excluded. For each epoch, the maximum head displacement was extracted, and the mean across all epochs were used as an index of head motion for each participant. The common artefact rejection threshold of 4000 fT in MEG studies of young children using SQUIDs (e.g., (Alho et al., 2023; W. He et al., 2015; Partanen et al., 2017)) was adjusted for the increased noise floor in OPM data using a factor of 13.78 (see (Safar et al., 2024) for further details). Participants were included in subsequent analyses if they had at least one minute of resting-state data remaining.

A linear regression identified a significant positive association between age and the number of epochs (*F*(1,63)=17.67, *p*<.001, *β*=0.47) and negative association between age and mean head motion (*F*(1,60)=39.75, *p*<.001, *β*=-0.65), respectively. To ensure these associations did not result in subsequent spurious developmental effects, the number of epochs was matched between the children and adults while minimizing the effect of head motion. For each adult, a number *N* was randomly drawn from the distribution of the number of epochs in the children. Then, the *N* epochs with the highest head motion were selected to be analyzed. Note that given the inherent relation between head motion and age, this effect was still significant after this procedure; however, unlike traditional SQUID-MEG systems, OPMs have been shown to be robust to head motion (Brookes et al., 2022).

The 90 cortical and subcortical regions of the automated anatomical labelling (AAL) atlas (Tzourio-Mazoyer et al., 2002) were reconstructed using a linearly constrained minimum variance (LCMV) beamformer. Forward solutions were computed for each source position using a single-shell head model of an age-appropriate template and assumed a dipole approximation of neural current (Nolte, 2003). For the adults, the International Consortium for Brain Mapping (ICBM) template in standard space (Fonov et al., 2009) was used. For the children, age-specific paediatric templates (Richards et al., 2016) were used, first warping the coordinates of each source in the AAL atlas from standard space to the templates using Advanced Normalization Tools (ANTs; version 2.4.3; (Avants et al., 2009)). The participants’ head digitisations collected during data acquisition and surface meshes of the age-appropriate templates were used to co-register the head models and source positions to the OPM data (R. M. Hill et al., 2020; Rhodes et al., 2023; Zetter et al., 2019). Covariance matrices for the beamformer were constructed across the continuous data and were regularized using the Tikhonov method with a regularization parameter of 2% (Tikhonov, 1943). To account for centre of the head biases, the neural activity index was applied by normalizing the beamformer output by the estimated noise (Van Veen et al., 1997).

### 2.4 Power spectra analyses

The reconstructed timeseries for each brain region were *z*-scored, and the power spectrum density (PSD) was computed using Welch’s method using window length of 1s with 50% overlap implemented in MATLAB (version R2021a; (The Mathworks Inc., 2018)); the resulting PSDs were averaged to obtain both whole-brain and regional absolute power spectra. Next, the absolute spectra were parameterized using the SpecParam algorithm (version 1.1.0; formally named Fitting Oscillations and One Over F (FOOOF)) implemented in Python (version 3.11.4), which models the spectra as a combination of aperiodic and periodic components (Donoghue, Haller, et al., 2020). The models were fitted between 2 and 40Hz, without modeling a bend, or knee, in the spectra. The periodic peaks modeled in the spectra were required to have bandwidths at least twice the frequency resolution (0.6Hz) of the spectra per the developer recommendations, with no other restrictions placed on the number or size of the peaks. The fits of the modelled spectra were visually inspected, and the correlation (*R*-squared) between the raw and fitted spectra and the error of the fitted model was used to evaluate model fit; no outliers (individuals with model fit metrics exceeding three standard deviations from the mean) were identified. For each model, the parameters of the aperiodic component (slope and offset) were extracted; the slope, alternatively called the exponent, captures the steepness of the exponential decay of a power spectrum, while the offset reflects a uniform shift of the spectrum across frequencies (Donoghue, Haller, et al., 2020). For the periodic component, the peak frequency and corresponding power were identified within bands of interest, selecting the modeled peak with the maximal power if multiple existed. The bands were derived by visually identifying peaks in periodic power spectra (computed by subtracting the aperiodic component from the absolute spectra) averaged across age group (see **Figure 1C** in the **Results**); widths were chosen to ensure the peaks were contained in all age groups. Three bands were identified: alpha (6-12Hz), low beta (13-20Hz), and high beta (21-25Hz).

**Figure 1:**
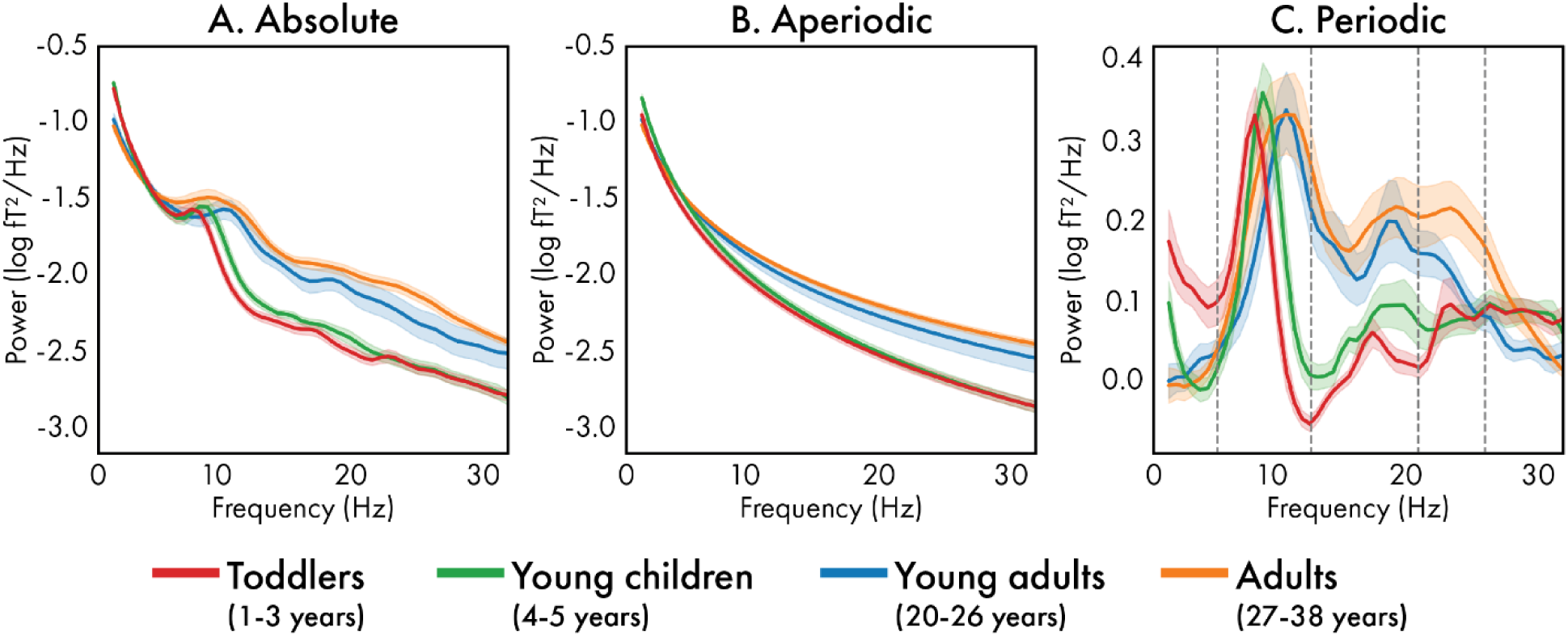
Whole-brain absolute (A), aperiodic (B), and periodic (C) power spectra averaged across the toddlers (1 – 3 years; red), young children (4 – 5 years; green), young adults (20 – 26 years; blue) and adults (27 – 38 years; orange).

### 2.5 Statistics

Regressions implemented in MATLAB (version R2021a; (The Mathworks Inc., 2018)) were used to test for associations between age and (a) sex, (b) the data quality measures (number of analyzed epochs, head motion), (c) model fit measures (error and *R*2), (d) aperiodic parameters (slope and offset), and (e) periodic parameters (presence or absence of a peak, peak frequency, and power for each frequency band). Logistic and linear regressions were used for binary and continuous variables, respectively, and standardized coefficients were reported as a measure of effect size. For sex, data quality, model fit, and whole-brain aperiodic and periodic measures, significance was held at *p*<0.05. For the regional aperiodic and periodic measures, *p*-values were corrected for multiple comparisons using the false discovery rate (FDR), holding significance at *q*<0.05. Associations between the aperiodic slope and offset were also investigated (see **Supplemental information** for further information).

There have been numerous demonstrations and discussions about the robustness of OPM-MEG to head motion given its wearable nature (Barry et al., 2019; Boto et al., 2018; R. M. Hill et al., 2019; Pedersen et al., 2022; Seedat et al., 2024; Seymour et al., 2021), and new paradigms have been designed where movement is in fact encouraged (Roberts et al., 2019). Nonetheless, we also demonstrate the lack of association between head motion and the measures of aperiodic and periodic activity in the young adults and adults, for whom age and head motion are not significantly correlated. These findings are presented in the **Supplemental information.**

## 3 Results

### 3.1 Participants

Sixty-nine participants were included in the analysis, after excluding four participants with too few trials remaining after quality control. Participant demographics and data quality measures are summarized by age group (toddlers, children, young adults, and adults) for descriptive purposes in **Table 1**, alongside regression statistics performed examining associations with continuous age. Age was not significantly associated with sex (*F*(1,67)=0.00, *p*=.977, *β*=0.01). After epoch selection, the number of epochs analyzed was not associated with age (*F*(1,67)=0.32, *p*=.572, *β*=0.07), but mean head motion was still found to significantly decrease with age (*F*(1,64)=33.47, *p*<.001, *β*=-0.60).

**Table 1:** Participant demographics, summarized by age group for descriptive purposes, with regression statistics examining associations with continuous age.

|  | Mean [SD] |  |  |  | Regression statistics |  |  |  |  |  |
| --- | --- | --- | --- | --- | --- | --- | --- | --- | --- | --- |
| | Toddlers | Young children | Young adults | Adults | *F* | *df* | *p*-value | *R*<sup>2</sup> | *B* [SE] ($\times 10^2$) | $\beta$ |
| *N* | 18 | 14 | 18 | 19 |  |  |  | – |  |  |
| **Age range (years)** | 1 - 3 | 4 - 5 | 20 - 26 | 27 - 38 |  |  |  | – |  |  |
| **Age (years)** | 3.0<br>[0.8] | 5.0<br>[0.5] | 24.0<br>[1.9] | 31.8<br>[3.5] |  |  |  | – |  |  |
| **Sex (F:M)** | 8:10 | 8:6 | 11:7 | 9:10 | 0.00 | (1,67) | .977 | 0.00 | 0.05<br>[1.92] | 0.01 |
| **# epochs** | 225.2<br>[68.7] | 254.1<br>[77.6] | 221.1<br>[73.6] | 246.6<br>[59.7] | 0.32 | (1,67) | .572 | 0.00 | 28.01<br>[66.92] | 0.07 |
| **Head motion (mm)** | 3.0<br>[2.0] | 3.8<br>[2.4] | 1.0<br>[0.7] | 1.0<br>[0.5] | 33.47 | (1,64) | <.001 | 0.34 | -9.35<br>[1.62] | -0.60 |

SD: standard deviation; *F*: *F*-statistic; *df*: degrees of freedom; *B*: coefficient; *SE*: standard error; $\beta$ : standardized coefficient; F: female; M: male

### 3.2 Power spectra analyses

Whole-brain and regional power spectra were modelled as a combination of aperiodic and periodic components (Donoghue, Haller, et al., 2020). The whole-brain absolute power spectra are presented in **Figure 1A** averaged by age group, alongside the modeled aperiodic power spectra (**Figure 1B**) and the periodic power spectra (**Figure 1C**). Descriptive statistics for the model fit metrics (*R*2 and error) for each age group are presented in **Supplemental Table 1**; there were no significant associations between age and the whole-brain model fit (*R*2: *F*(1,67)=2.16, *p*=.146, *β*=-0.18; error: *F*(1,67)=0.00, *p*=.997, *β*=0.00).

All statistics for the whole-brain aperiodic and periodic measures are presented in **Table 2**. The slope of the whole-brain aperiodic component was found to significantly decrease with age (**Figure 2A**; *F*(1,63)=40.94, *p*<.001, *β*=-0.62); this association was also significant in all cortical and subcortical brain regions (**Figure 2B**), with the strongest effects observed in the left pre- and post-central gyri, the left superior and inferior parietal gyri, and the bilateral cuneus and precuneus. The offset of the whole-brain (**Figure 2C**; *F*(1,63)=24.17, *p*<.01, *β*=-0.51) and regional (**Figure 2D**) aperiodic components also significantly decreased with age, with the regional effects significant except for the orbital frontal cortices and temporal poles. Decreases in the aperiodic offset followed a similar pattern as the aperiodic slope. Associations between the slope and offset are presented in the **Supplemental information**.

**Figure 2:**
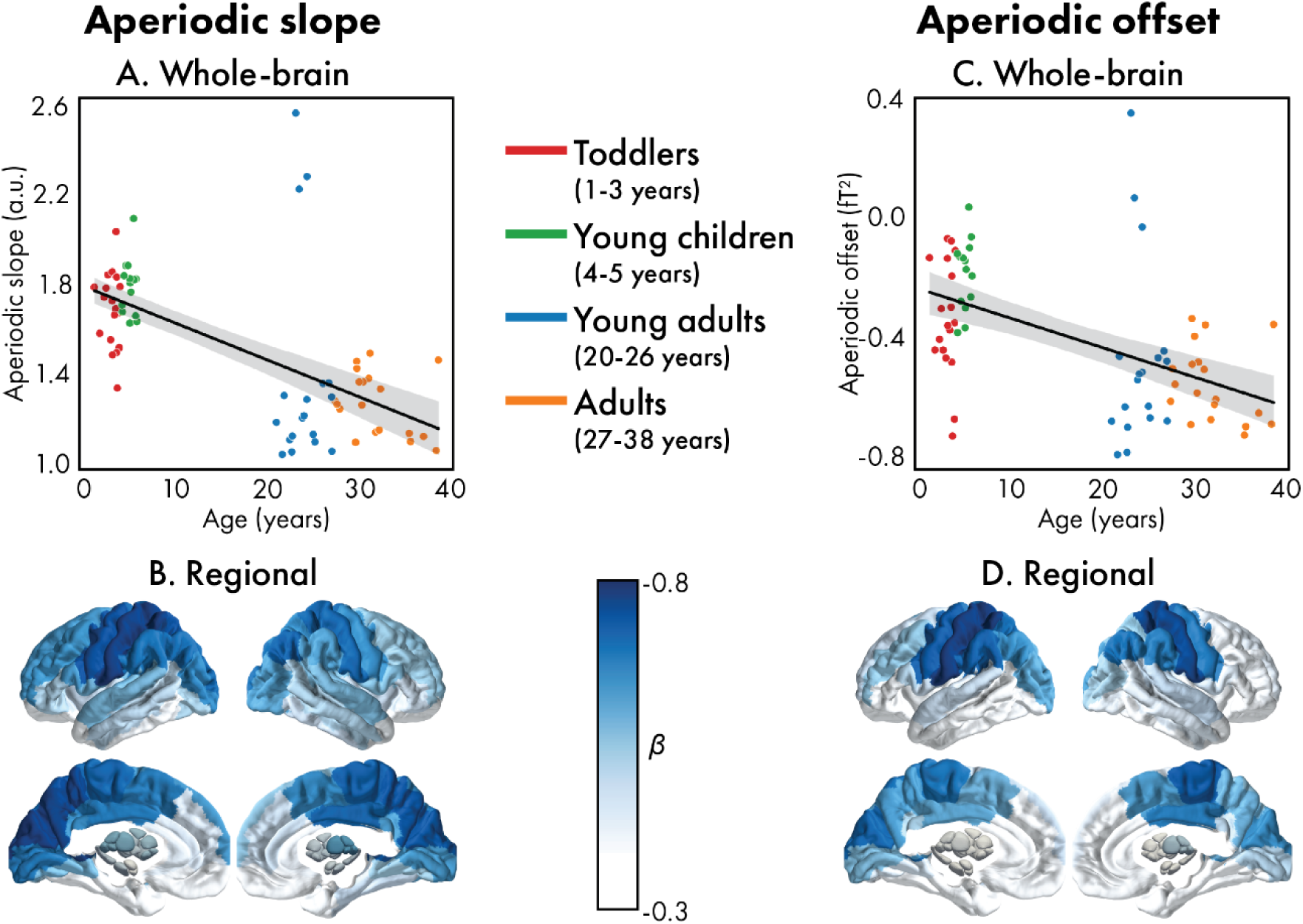
Associations between the **aperiodic** slope (left: A and B) and offset (right: C and D) for both the whole-brain (top: A and C) and regional (bottom: B and D) power spectra. For the whole-brain spectra, values are coloured according to age group (red: toddlers, green: young children, blue: young adults, orange: adults). For the regional spectra, the standardized coefficients (*β*) for significant (corrected) brain regions are presented.

**Table 2:**
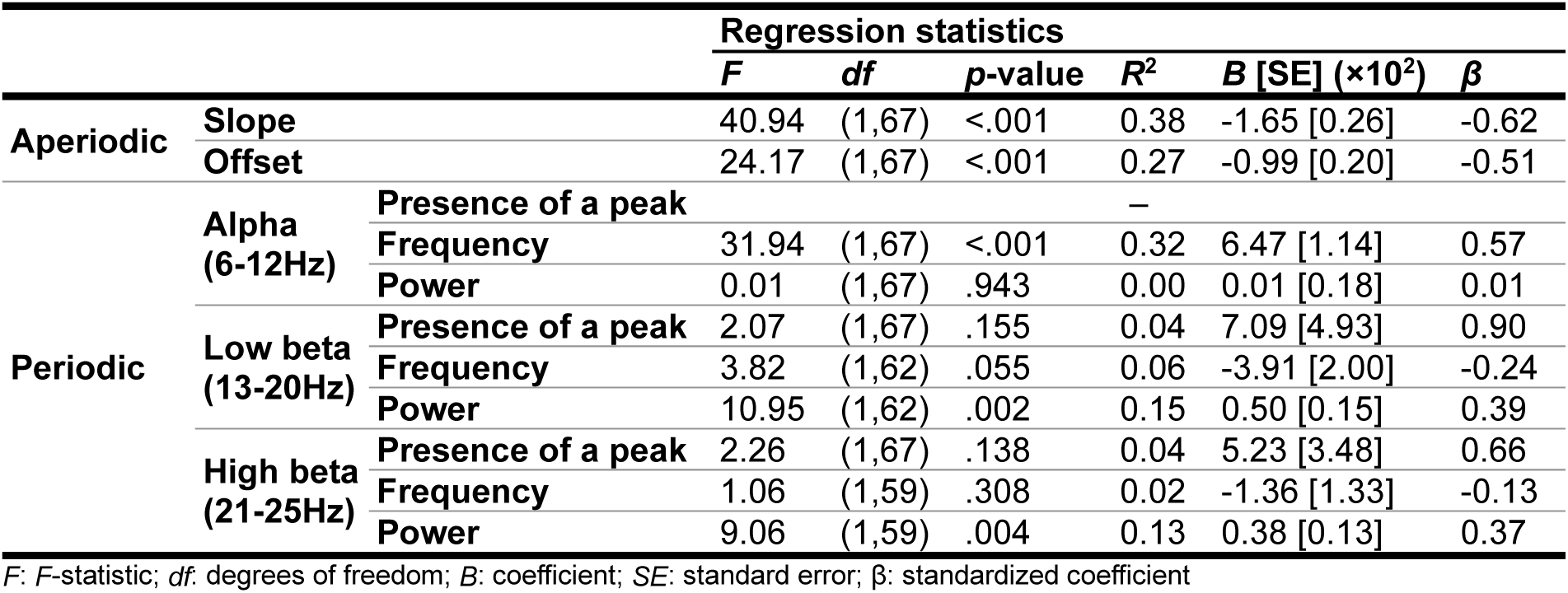
Regression statistics examining associations between age and the aperiodic, and periodic parameters.

The whole-brain models identified peaks in all participants in the alpha frequency band. In both low and high beta, peaks were identified in 93% and 88% of participants, respectively, and the presence of a peak was not associated with age (low beta: *F*(1,67)=2.07, *p*=.155, *β*=0.90; high beta: *F*(1,67)=2.26, *p*=.138, *β*=0.66). The percentage of participants with peaks on a regional level is presented in **Supplemental Figure 1**; in all frequency bands, there was no significant associations with age in any brain region.

In alpha (6-12Hz; **Figure 3A**), whole-brain peak frequency increased with age (**Table 2**; *F*(1,67)=31.94, *p*<.001, *β*=0.57), with all brain regions also showing this pattern; largest effects were observed bilateral parietal regions, particularly the pre- and post-central gyri, medial occipital regions, subcortical regions, and the posterior and median cingulate gyri. Age-related changes in peak alpha power was not significant at a whole-brain (*F*(1,67)=0.01, *p*=.943, *β*=0.01) or regional level. In low beta (13-20Hz; **Figure 3B**), whole-brain peak frequency was not associated with age (*F*(1,62)=3.82, *p*=.055, *β*=0.39), nor in any individual brain region. Low beta power increased with age across the whole brain (*F*(1,62)=10.95, *p*=.002, *β*=0.39) as well as in 74 of the 90 brain regions, with the largest effects occurring in bilateral temporal regions, bilateral occipital regions, particularly the calcarine cortices, and the left supramarginal gyrus. In high beta (21-30Hz; **Figure 3C**), while whole-brain peak frequency did not significantly decrease with age (*F*(1,59)=1.06, *p*=.308, *β*=-0.13), this effect was significant in the opercular part of the right inferior frontal gyrus, the right middle temporal gyrus, and the right Heschl’s gyrus. On the other hand, high beta power increased with age globally (*F*(1,59)=9.06, *p*=.004, *β*=0.37) as well as regionally in 65 regions throughout the brain, with strong effects in the bilateral frontal lobe, particularly in the right hemisphere, the right temporal pole, bilateral subcortical regions, and the anterior cingulate.

**Figure 3:**
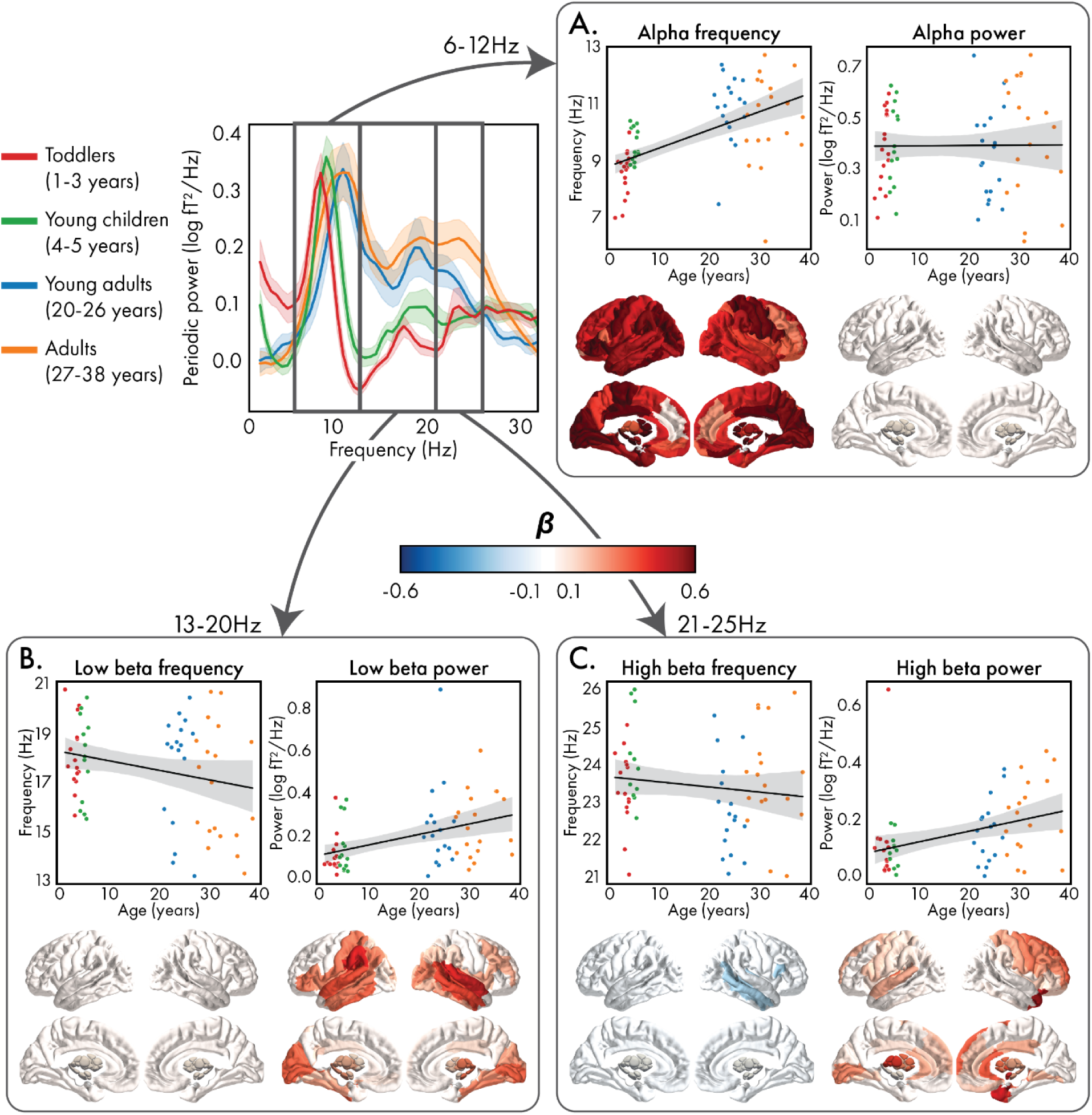
Associations between age and the parameters of the **periodic** power spectra (center) in the alpha (A), low beta (B), and high beta (C) frequency bands. In each panel, peak frequency (left) and corresponding power (right) are shown for the whole-brain (top) and regional (bottom) power spectra. For the whole-brain measures, values are coloured according to age group (red: toddlers, green: young children, blue: young adults, orange: adults). For the regional spectra, the standardized coefficients (*β*) for significant (corrected) brain regions are presented.

## 4 Discussion

This study is the first to use OPMs, a new, wearable MEG technology, to measure brain function in very young children, and the first to examine changes in periodic and aperiodic components of neural signals between very early childhood (1 – 5 years of age) and adulthood (20 – 38 years of age). Consistent with prior EEG studies, we found significant developmental effects of the aperiodic (slope and offset) and periodic (peak frequency and power) features of brain function using OPM-MEG. Additionally, we are the first to show that these changes varied across distinct brain regions. This work not only adds to the growing body of work highlighting the advantages of using OPMs ((Pedersen et al., 2022; Rhodes et al., 2023; Safar et al., 2024; Wittevrongel et al., 2021), for a review see (Brookes et al., 2022)), especially for studying development (Boto et al., 2022; Feys & De Tiège, 2024; R. M. Hill et al., 2019), but also contributes novel insights into the spatial pattern of the effect sizes of neurophysiological changes with age.

While EEG and SQUID-MEG can capture the neurophysiological measurements required to fully characterize the development of aperiodic and periodic activity in the brain, they present several important limitations. While EEG is wearable and offers good temporal resolution, the conductive properties of the skull limit its spatial resolution (Hari & Salmelin, 2012), signals are affected by skull thickness (Hoekema et al., 2003), and signals are susceptible to muscle artefacts that occur during head movement (Boto et al., 2018). While SQUID-MEG systems have good temporal and spatial resolution, the need for cryogenic cooling requires the systems to be fixed and have a “one size fits all” helmet, typically designed to fit the average adult head size, which can lead to inhomogeneous coverage and a reduction in signal for smaller compared to larger head sizes; SQUID-MEG signals are also susceptible to head motion (Gross et al., 2013).

Given the age-related associations with skull thickness, head size, and head motion, these issues have posed challenges for developmental EEG and SQUID-MEG studies that have spanned early childhood and adulthood (Brookes et al., 2022). OPM-MEG systems address these issues, offering the spatial resolution of SQUID-MEG with the wearable nature of EEG, making it particularly suitable for studying the development of brain function across the life span. Our work is the first to leverage these advantages to examine the development of aperiodic and periodic activity between childhood and adulthood. We show that well-established developmental patterns, such as the increase in peak alpha frequency, can be replicated by this technology, while also providing new insight into brain development that was only possible using OPM-MEG.

### 4.1 Developmental changes in aperiodic signals

We found that both aperiodic slope and offset significantly decreased between toddlerhood and adulthood. By examining changes between early toddlerhood (1 – 5 years of age) and adulthood (20 – 38 years of age), we extend previous developmental EEG work that found similar decreases across smaller age ranges (Cellier et al., 2021; Favaro et al., 2023; A. T. Hill et al., 2022; McSweeney et al., 2021, 2023; Schaworonkow & Voytek, 2021; Tröndle et al., 2022). We also provide important insights into how the effect sizes of these decreases differ across the brain. While previous work has only been able to characterize developmental patterns with broad, lobe-level specificity due to the poorer spatial resolution of EEG (Cellier et al., 2021; Favaro et al., 2023; A. T. Hill et al., 2022; Schaworonkow & Voytek, 2021), the spatial resolution of OPM-MEG is comparatively much improved, and thus we were able to determine the strength of these changes within individual brain regions.

While the precise neurobiological mechanisms underpinning the aperiodic slope remain unclear, there is evidence to support an association with E/I balance (Gao et al., 2017). Excitatory and inhibitory activity is characterized by faster and slower synaptic currents, respectively, which corresponds to a difference in spectral decay (Gao et al., 2017): increased excitation relative to inhibition results in a flatter power spectrum, while increased inhibition relative to excitation results in a steeper spectrum. The link between E/I balance and the spectral slope has been supported by studies examining changes in the aperiodic slope under pharmacological interventions known to modify inhibitory and excitatory neural activity (Colombo et al., 2019; Gao et al., 2017; Lendner et al., 2020; Medel et al., 2023; Waschke et al., 2021). Furthermore, increases in excitation relative to inhibition can temporally decorrelate spiking in neuronal populations causing neuronal “noise” (Voytek & Knight, 2015), which has been shown to relate to flatter spectral slopes (Pozzorini et al., 2013; Usher et al., 1995). Thus, the observed age-related flattening of the spectral slope may reflect maturational changes in E/I balance, specifically a shift towards excitation with increased age, consistent with the existing developmental EEG work (Cellier et al., 2021; Favaro et al., 2023; A. T. Hill et al., 2022; McSweeney et al., 2021, 2023; Schaworonkow & Voytek, 2021; Tröndle et al., 2022).

This conclusion is supported by evidence of protracted changes towards increasingly balanced E/I throughout development and into adulthood that are hypothesized to support plasticity (Larsen et al., 2022; Perica et al., 2022), and maturational increases of neural noise that is thought to reflect the expansion of the brain’s “dynamic repertoire” of functional network configurations (McIntosh et al., 2010).

Shifts in the aperiodic offset can be interpreted as changes in broadband power (Donoghue, Haller, et al., 2020). As such, the aperiodic offset is thought to reflect the overall spiking activity of neuronal populations (Manning et al., 2009; Miller et al., 2009, 2014), and it has been hypothesized that the synaptic pruning, which causes reductions in cortical gray matter throughout childhood, is driving the observed decreases in offset (Cellier et al., 2021; Favaro et al., 2023; A. T. Hill et al., 2022; McSweeney et al., 2023; Tröndle et al., 2022). However, it has not yet been possible to exclude age-related confounds as the driver of this effect. All published developmental studies to date have used EEG, and increases in skull thickness, which occurs across development, results in decreases in the amplitudes of EEG signals (Hoekema et al., 2003). Given that the aperiodic slope is proportional to broadband power (Donoghue, Haller, et al., 2020), the age-related shifts in the broadband power may simply reflect developmental changes other than the brain (A. T. Hill et al., 2022; McSweeney et al., 2023). On the other hand, magnetic fields can pass through the skull without distortion (Okada et al., 1999), and thus our OPM-MEG study can conclude that reductions in aperiodic offset with age are genuine. It is well established that cortical gray matter rapidly expands during early childhood, followed by a gray matter reduction in childhood, with protracted changes extending through adolescence (Norbom et al., 2021). The decreases in cortical gray matter can be attributed, in part, to the synaptic pruning and increased myelination that occurs during childhood and adolescence to support the development of efficient neural circuitry (Gogtay et al., 2004; Huttenlocher & Dabholkar, 1997; Paolicelli et al., 2011).

Thus, the age-related reductions in global spectral power reported here, and elsewhere (Cellier et al., 2021; Favaro et al., 2023; Gómez et al., 2017; A. T. Hill et al., 2022; McSweeney et al., 2021, 2023; Miskovic et al., 2015; Schaworonkow & Voytek, 2021; Tröndle et al., 2022), which have been shown to parallel the changes in cortical thickness (Whitford et al., 2007), may be due to synaptic pruning and apparent reduction of grey matter.

However, the aperiodic slope and offset are highly correlated (Donoghue, Haller, et al., 2020). In our study, analyses revealed that the whole-brain aperiodic slope and offset were strongly associated (see the **Supplemental information**), which is consistent with other examinations of this relation (Favaro et al., 2023; McSweeney et al., 2021, 2023; Tröndle et al., 2022). Since we were able to measure the aperiodic components on a regional level, we were also able to demonstrate that there was a high degree of correlation between slope and offset within each brain region, as well as with respect to the patterns of age-related changes across the brain (**Supplemental information**). While changes in the offset can occur in the absence of a change in the slope (e.g., a vertical shift in the spectrum), a flatter or steeper slope necessitates a co-occurring shift in the offset, if the spectrum is rotated around a non-zero frequency (Donoghue, Haller, et al., 2020; Ostlund et al., 2022). However, the developmental trajectories of the two components were shown to be distinct in the first months of life across different vigilance stages (Favaro et al., 2023), suggesting that the developmental changes in slope and offset are due to distinct neurophysiological processes (e.g., E/I balance and synaptic pruning, respectively).

The sequence of the development of brain function follows the hierarchical organization of the brain, with the lower-order, sensorimotor cortices maturing by early childhood, while the maturation of higher-order, association cortices is protracted, extending throughout adolescence (e.g., (Dong et al., 2021; Pines et al., 2022), see (Grayson & Fair, 2017; Norbom et al., 2021; Sydnor et al., 2021) for reviews). The pattern of the effect sizes for the age-related changes in the aperiodic components follows a similar pattern: unimodal regions, such as the motor and visual cortices, show larger effects, while higher-order regions, such as the frontal cortices, show smaller effects. This broadly aligns with the existing EEG literature, where studies examining the topography of developmental changes reported steeper decreases in aperiodic activity in parietal-occipital compared to frontal electrodes (Favaro et al., 2023; A. T. Hill et al., 2022; Schaworonkow & Voytek, 2021). Previous work has provided evidence that aperiodic activity changes non-linearly between early-to-middle childhood, with activity following an inverted-U shaped trajectory, peaking at approximately seven years of age (McSweeney et al., 2023). We propose that our findings coincide with this quadratic developmental model of aperiodic activity and provide evidence that the sequence of development across the brain follows the sensorimotor-association axis. Specifically, if the aperiodic activity of lower-order regions has already peaked, or is close to peaking, during early childhood, changes between childhood and adulthood would be steep. On the other hand, if the development of higher-order regions is protracted, not reaching peak activity until adolescence, changes between childhood and adulthood would be comparatively flatter. While this pattern of non-linear and hierarchical development aligns with the broader functional neuroimaging literature (Hunt et al., 2019; Sydnor et al., 2021), conclusions can not be drawn firmly without examining changes in aperiodic activity across childhood, adolescence, and adulthood.

### 4.2 Developmental changes in periodic signals

Our finding of developmental increases of peak alpha frequency is well-documented (e.g., (Cellier et al., 2021; Cragg et al., 2011; Gómez et al., 2017; A. T. Hill et al., 2022; Marshall et al., 2002; McSweeney et al., 2023; Miskovic et al., 2015; Rodríguez-Martínez et al., 2017; Tröndle et al., 2022)), and is thought to coincide with increases in the speed of neural communication to facilitate neurocognitive development (Cellier et al., 2021; McSweeney et al., 2023; Segalowitz et al., 2010; Tröndle et al., 2022). This work shows that OPM-MEG replicates this finding and provides insight into the precise spatial patterns of this maturational change. All examined brain regions showed significant increases in peak alpha frequency with age, consistent with a previous report that, aside from increasing frequency with age, the topographies of alpha spectral power were similar between children and adults (Rodríguez-Martínez et al., 2017). We found no changes in alpha power with age aligning with some studies (e.g., (Clarke et al., 2001)), but not others, with reports of significant age-related increases (e.g., (Tröndle et al., 2022)) and decreases (e.g., (Whitford et al., 2007)). A recent, well-powered study reported increases in alpha power between 5 and 22 years of age after adjusting for aperiodic activity and suggested that the conflicting reports were due to the computation of alpha power across canonical frequency bands and unadjusted alpha power (Tröndle et al., 2022). Our methodology was similar to that of Tröndle and colleagues and thus also addressed these concerns, however we were not able to identify an increase in alpha power. While this could be due to different age ranges across the two studies, we also note that different resting-state conditions were used (*Inscapes* in the current study, compared to eyes-closed resting-state). Alpha power in eyes open versus eyes closed resting-states was found to show diverging developmental trajectories (A. T. Hill et al., 2022; McSweeney et al., 2023), and thus we hypothesize that the lack of reported maturation of alpha power may be due to the use of an alternative, eyes open resting-state paradigm and differing age ranges.

Two peaks were identified in the beta frequency band: low (13 – 20Hz) and high (21 – 25Hz). Two beta bands have been identified in resting-state EEG (Rosanova et al., 2009) and MEG (Capilla et al., 2022; Mahjoory et al., 2020) data, and they have been shown to be generated by different mechanisms (Cannon et al., 2014); however, we are one of the first to identify distinct developmental patterns between them. In this study, we found that both beta bands showed increases in power with age. Increased beta power between childhood and adulthood is consistent with previous work (Gómez et al., 2017; Heinrichs-Graham et al., 2018; Schäfer et al., 2014), with evidence to suggest these differences do not emerge until adolescence (A. T. Hill et al., 2022), and are thought to support the development of sensorimotor and cognitive control (Engel & Fries, 2010). High beta power has also been differentially associated with frontal regions (Capilla et al., 2022; Ferrarelli et al., 2012; Rosanova et al., 2009). For example, transcranial magnetic stimulation (TMS) was found to evoke low beta oscillations in parietal and perirolandic regions, and high beta oscillations in the frontal cortex (Ferrarelli et al., 2012; Rosanova et al., 2009). Relatedly, a data-driven atlas of resting-state oscillations constructed using MEG found that low beta oscillations were specific to lateral occipital-parietal regions, while high beta oscillations were specific to motor and prefrontal cortices. This aligns with our finding that age-related changes in high-beta power have the strongest effects in the frontal lobe and is further supported by the increasing changes between the young and slightly older adults, as brain maturation, particularly in the frontal and association cortices continues throughout young adulthood. Interestingly, alongside occipital and parietal regions, temporal regions also showed strong maturational effects of low-beta power. Given the importance of the temporal lobes in social cognition (Olson et al., 2013), this may reflect increases in these skills with age, and warrants further investigation. On the other hand, peak frequency only decreased with age in the high beta band. To our knowledge, only one study from 1999 has investigated the maturational effects of peak frequency separately for low and high beta, and they also found significant decreases in high beta frequency with age, while low beta showed no change (Dustman et al., 1999). Future work is needed to understand the importance of these diverging patterns.

### 4.3 Limitations

This study has several limitations. While we were able to characterize the changes between very young children and adults, our sample lacked participants between 6-19 years of age, which is an important period of neurodevelopment. This study leveraged two retrospective OPM-MEG datasets, one of adults which was collected to replicate SQUID-MEG data (Safar et al., 2024), and the other is a sample of typically developing toddlers and young children being collected to study the neurophysiological differences of autism early in life (study is ongoing). OPM-MEG data from school-age children and adolescents were not available, and thus we were unable to include this age range in our analysis. Furthermore, nonlinear maturational trajectories in brain function are prevalent in the neuroimaging literature (Hunt et al., 2019; Sydnor et al., 2021), however, we cannot provide insight into the nature of changes between toddlerhood/early childhood and early adulthood without examining the full spectrum of childhood and adolescence. Our findings must be interpreted in the absence of school-age children and adolescents, and future studies should examine their replicability across development. Our study was also cross-sectional – the generalization of this work to longitudinal changes in the brain is important for future research. While we hypothesize that age-related changes in aperiodic and periodic activity are important for development, future work examining these changes alongside changes in cognition and behaviour will be important for contextualizing these findings. Our findings in deep brain structures should also be interpreted with caution, especially with respect to the measures of periodic activity where effect sizes were large. While the customizable helmets in OPM-MEG addresses the concern of using one-size-fits-all helmets in SQUID-MEG to study development, the smaller head circumferences in children relative to adults means the sensors are relatively closer to subcortical regions in the children compared to adults, which would increase signal-to-noise.

### 4.4 Conclusions

This is the first study to use OPM-MEG to investigate how aperiodic and periodic activity develops in very young children and early adulthood. We found age-related changes in spectral features measured using OPM-MEG that were consistent with the existing literature but leveraged MEG’s spatial resolution to report, for the first time, that the effect sizes of these changes differ throughout the brain. Click or tap here to enter text.Our work demonstrates the utility of OPM-MEG for studying developmental changes in brain function, especially changes over the early years of life that cannot be reliably measured using traditional adult-sized SQUID-MEG and lays the foundation for future studies to examine aperiodic and periodic activity across the life span and their role in developmental disorders.

## 5 Acknowledgements

Funding was provided by the Canadian Institutes of Health Research (PJT-178370) and the Simons Foundation Autism Research Initiative (SFARI; 2021 Human Cognitive and Behavioural Science award). We would like to thank all individuals and their families who participated in this study and those who helped with data collection.

## 6 Author contributions

Conceptualization: M. M. Vandewouw, M. J. Taylor; Formal analysis: M. M. Vandewouw; Funding acquisition: M. J. Taylor; Investigation: J. Sato, N. Rhodes, K. Safar; Methodology: M. M. Vandewouw; Project administration: J. Sato, M. J. Taylor; Resources: M. J. Taylor; Software: M. M. Vandewouw; Supervision: M. J. Taylor; Visualization: M. M. Vandewouw; Roles/Writing - original draft: M. M. Vandewouw, M. J. Taylor; and Writing - review & editing: M. M. Vandewouw, J. Sato, K. Safar, M. J. Taylor.

## Abbreviations

(OPM): optically pumped magnetometer
(MEG): magnetoencephalography
(EEG): electroencephalography
(E/I): excitatory-inhibitory
(SQUID): superconducting quantum interference device
(PSD): power spectrum density
(FOOOF): Fitting Oscillations and One Over F
(SD): standard deviation
(F): *F*-statistic
(df): degrees of freedom
(B): regression coefficient
(SE): standard error
(β): standardized regression coefficient

## 9 Supplemental information

### 9.1 Model fit metrics

**Supplemental Table 1:** Model fit metrics, summarized by age group for descriptive purposes, with regression statistics examining associations with continuous age.

|  | Mean [SD] |  |  |  | Regression statistics |  |  |  |  |  |
| --- | --- | --- | --- | --- | --- | --- | --- | --- | --- | --- |
|  | Toddlers | Young children | Young adults | Adults | F | df | p-value | R<sup>2</sup> | B [SE] (×10<sup>2</sup>) | β |
| R<sup>2</sup> | 0.995<br>[0.005] | 0.998<br>[0.001] | 0.996<br>[0.002] | 0.995<br>[0.003] | 2.16 | (1,67) | .146 | 0.03 | -0.00<br>[0.00] | -0.18 |
| Error | 0.023<br>[0.011] | 0.016<br>[0.005] | 0.019<br>[0.076] | 0.021<br>[0.007] | 0.00 | (1,67) | .997 | 0.00 | -0.00<br>[0.01] | 0.00 |

SD: standard deviation; F: F-statistic; df: degrees of freedom; B: coefficient; SE: standard error; β: standardized coefficient

### 9.2 Regional presence of peaks

**Supplemental Figure 1:**
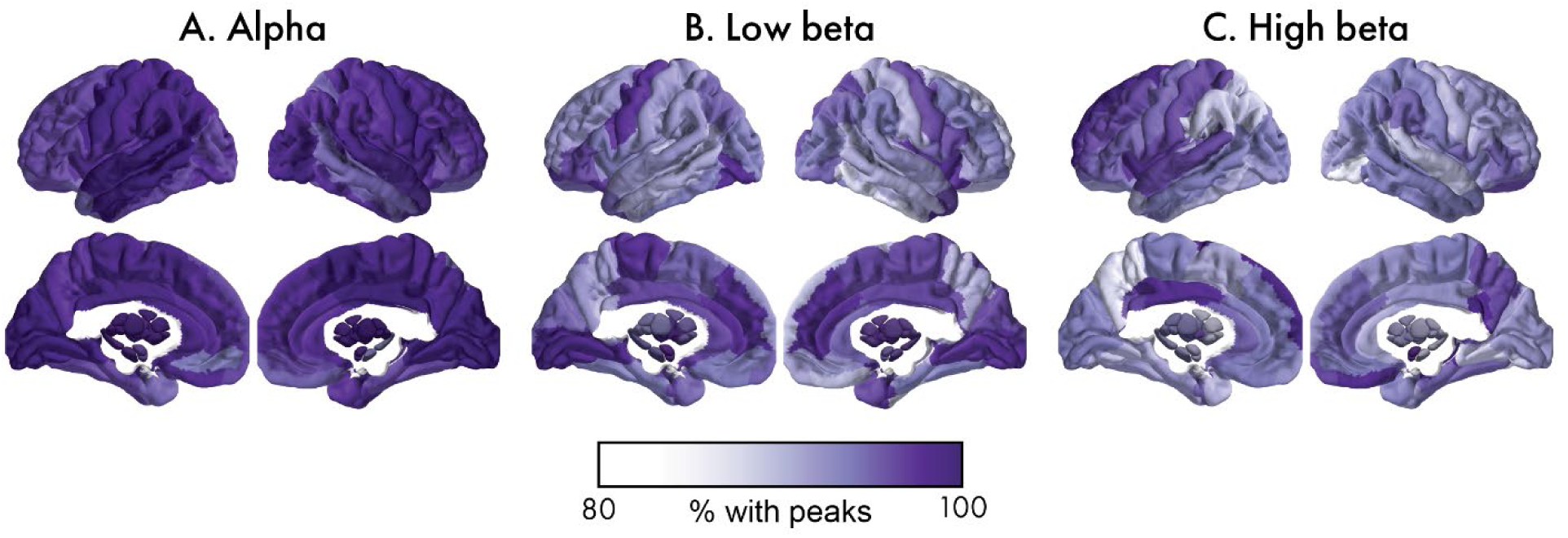
Percent of participants with detectable peaks for each brain region for alpha (A), low beta (B), and high beta (C).

### 9.3 Associations between aperiodic slope and offset

To examine the association between whole-brain aperiodic slope and offset, we performed a linear mixed effects regression with offset as the dependent variable, slope as the independent variable, and participant-specific random intercepts. A significant positive association was found (*F*(1,67)=443.3, *p*<.001, *β*=0.93). We also applied these models to each brain region, FDR-correcting for multiple comparisons. Significant positive associations were observed in each of the 90 brain regions, with effect sizes (standardized coefficients, equivalent to Pearson correlation coefficients for these models) from 0.75 to 0.95. To examine whether age-related changes in the aperiodic slope and offset were also associated across the brain, a regression was performed between the regional effect sizes (**Figure 2B** and **2D**). Again, a significant association was found (*F*(1,88)=805.1, *p*<.001, *β*=0.95).

### 9.4 Associations with head motion

Within the young adult and adult participants, there was no significant association between head motion and age (*F*(1,33)=0.16, *p*=.694, *β*=-0.08). In this sample, there were no significant associations between head motion and the whole-brain aperiodic and periodic features (**Supplemental Table 2**), nor were there any significant associations at the regional level. On top of the established robustness of OPM-MEG to head motion, the lack of association between head motion and aperiodic/periodic activity in the adult samples buttresses our conclusions that our findings are not being driven by age-related differences in head motion.

**Supplemental Table 2:** Regression statistics examining associations between head motion and the aperiodic, and periodic parameters in the young adult and adult samples, which show no association between head motion and age.

|  |  |  | Regression statistics |  |  |  |  |  |
| --- | --- | --- | --- | --- | --- | --- | --- | --- |
| | | | *F* | *df* | *p*-value | *R*<sup>2</sup> | *B* [SE] ($\times 10^2$) | $\beta$ |
| Aperiodic | Slope |  | 1.33 | (1, 35) | .257 | 0.04 | -1.34 [1.16] | -0.19 |
|  | Offset |  | 1.10 | (1, 35) | .301 | 0.03 | -0.84 [0.80] | -0.17 |
| Periodic | Alpha (6-12Hz) | Presence of a peak | - |  |  |  |  |  |
|  |  | Frequency | 0.06 | (1, 35) | .803 | 0.00 | -1.22 [4.84] | -0.04 |
|  |  | Power | 0.00 | (1, 35) | .950 | 0.00 | 0.04 [0.72] | 0.01 |
|  | Low beta (13-20Hz) | Presence of a peak | 0.05 | (1, 35) | .830 | 0.00 | 4.92 [22.69] | 0.24 |
|  |  | Frequency | 1.82 | (1, 34) | .186 | 0.05 | -10.66 [7.90] | -0.22 |
|  |  | Power | 0.15 | (1, 34) | .693 | 0.00 | 0.25 [0.62] | 0.07 |
|  | High beta (21-25Hz) | Presence of a peak | 0.11 | (1, 35) | .738 | 0.00 | -4.89 [14.51] | -0.24 |
|  |  | Frequency | 0.80 | (1, 33) | .376 | 0.02 | 4.26 [4.76] | 0.15 |
|  |  | Power | 4.10 | (1, 33) | .051 | 0.11 | 0.84 [0.42] | 0.33 |

*F*: *F*-statistic; *df*: degrees of freedom; *B*: coefficient; *SE*: standard error; $\beta$ : standardized coefficient

## References

Alho, J., Samuelsson, J. G., Khan, S., Mamashli, F., Bharadwaj, H., Losh, A., McGuiggan, N. M., Graham, S., Nayal, Z., Perrachione, T. K., Joseph, R. M., Stoodley, C. J., Hämäläinen, M. S., & Kenet, T. (2023). Both stronger and weaker cerebro-cerebellar functional connectivity patterns during processing of spoken sentences in autism spectrum disorder. Human Brain Mapping, 44(17), 5810–5827. 10.1002/HBM.26478

Avants, B. B., Tustison, N., & Song, G. (2009). Advanced normalization tools (ANTS). The Insight Journal, 2(365), 1–35. 10.54294/uvnhin

Baillet, S. (2017). Magnetoencephalography for brain electrophysiology and imaging. Nature Neuroscience 2017 *20*:3, *20*(3), 327–339. 10.1038/NN.4504

Barry, D. N., Tierney, T. M., Holmes, N., Boto, E., Roberts, G., Leggett, J., Bowtell, R., Brookes, M. J., Barnes, G. R., & Maguire, E. A. (2019). Imaging the human hippocampus with optically-pumped magnetoencephalography. NeuroImage, 203, 116192. 10.1016/J.NEUROIMAGE.2019.116192

Boto, E., Holmes, N., Leggett, J., Roberts, G., Shah, V., Meyer, S. S., Muñoz, L. D., Mullinger, K. J., Tierney, T. M., Bestmann, S., Barnes, G. R., Bowtell, R., & Brookes, M. J. (2018). Moving magnetoencephalography towards real-world applications with a wearable system. Nature 2018 555:7698, 555(7698), 657–661. 10.1038/nature26147

Boto, E., Shah, V., Hill, R. M., Rhodes, N., Osborne, J., Doyle, C., Holmes, N., Rea, M., Leggett, J., Bowtell, R., & Brookes, M. J. (2022). Triaxial detection of the neuromagnetic field using optically-pumped magnetometry: feasibility and application in children. NeuroImage, 252, 119027. 10.1016/J.NEUROIMAGE.2022.119027

Brookes, M. J., Leggett, J., Rea, M., Hill, R. M., Holmes, N., Boto, E., & Bowtell, R. (2022). Magnetoencephalography with optically pumped magnetometers (OPM-MEG): the next generation of functional neuroimaging. Trends in Neurosciences, 45(8), 621–634. 10.1016/j.tins.2022.05.008

Buzsáki, G., & Draguhn, A. (2004). Neuronal olscillations in cortical networks. Science, 304(5679), 1926–1929. 10.1126/SCIENCE.1099745/ASSET/F6069D2C-D3F3-4DAD-BB7A-4410FF87F651/ASSETS/GRAPHIC/ZSE0250426280003.JPEG

Cannon, J., Mccarthy, M. M., Lee, S., Lee, J., Börgers, C., Whittington, M. A., & Kopell, N. (2014). Neurosystems: brain rhythms and cognitive processing. European Journal of Neuroscience, 39(5), 705–719. 10.1111/EJN.12453

Capilla, A., Arana, L., García-Huéscar, M., Melcón, M., Gross, J., & Campo, P. (2022). The natural frequencies of the resting human brain: An MEG-based atlas. NeuroImage, 258, 119373. 10.1016/J.NEUROIMAGE.2022.119373

Cellier, D., Riddle, J., Petersen, I., & Hwang, K. (2021). The development of theta and alpha neural oscillations from ages 3 to 24 years. Developmental Cognitive Neuroscience, 50, 100969. 10.1016/J.DCN.2021.100969

Cesnaite, E., Steinfath, P., Jamshidi Idaji, M., Stephani, T., Kumral, D., Haufe, S., Sander, C., Hensch, T., Hegerl, U., Riedel-Heller, S., Röhr, S., Schroeter, M. L., Witte, A. V., Villringer, A., & Nikulin, V. V. (2023). Alterations in rhythmic and non-rhythmic resting-state EEG activity and their link to cognition in older age. NeuroImage, 268, 119810. 10.1016/J.NEUROIMAGE.2022.119810

Clarke, A. R., Barry, R. J., McCarthy, R., & Selikowitz, M. (2001). Age and sex effects in the EEG: development of the normal child. Clinical Neurophysiology, 112(5), 806– 814. 10.1016/S1388-2457(01)00488-6

Colombo, M. A., Napolitani, M., Boly, M., Gosseries, O., Casarotto, S., Rosanova, M., Brichant, J. F., Boveroux, P., Rex, S., Laureys, S., Massimini, M., Chieregato, A., & Sarasso, S. (2019). The spectral exponent of the resting EEG indexes the presence of consciousness during unresponsiveness induced by propofol, xenon, and ketamine. NeuroImage, 189, 631–644. 10.1016/J.NEUROIMAGE.2019.01.024

Cragg, L., Kovacevic, N., McIntosh, A. R., Poulsen, C., Martinu, K., Leonard, G., & Paus, T. (2011). Maturation of EEG power spectra in early adolescence: a longitudinal study. Developmental Science, 14(5), 935–943. 10.1111/J.1467-7687.2010.01031.X

Dong, H. M., Margulies, D. S., Zuo, X. N., & Holmes, A. J. (2021). Shifting gradients of macroscale cortical organization mark the transition from childhood to adolescence. Proceedings of the National Academy of Sciences of the United States of America, 118(28), e2024448118. 10.1073/PNAS.2024448118/SUPPL_FILE/PNAS.2024448118.SD02.TXT

Donoghue, T., Dominguez, J., & Voytek, B. (2020). Electrophysiological Frequency Band Ratio Measures Conflate Periodic and Aperiodic Neural Activity. ENeuro, 7(6). 10.1523/ENEURO.0192-20.2020

Donoghue, T., Haller, M., Peterson, E. J., Varma, P., Sebastian, P., Gao, R., Noto, T., Lara, A. H., Wallis, J. D., Knight, R. T., Shestyuk, A., & Voytek, B. (2020). Parameterizing neural power spectra into periodic and aperiodic components. Nature Neuroscience 2020 *23*:12, *23*(12), 1655–1665. 10.1038/s41593-020-00744-x

Dustman, R. E., Shearer, D. E., & Emmerson, R. Y. (1999). Life-span changes in EEG spectral amplitude, amplitude variability and mean frequency. Clinical Neurophysiology, 110(8), 1399–1409. 10.1016/S1388-2457(99)00102-9

Ebersole, J. S., & Pedley, T. A. (2003). Current practice of clinical electroencephalography, 3rd edn. European Journal of Neurology, 10(5), 604–605. 10.1046/J.1468-1331.2003.00643.X

Engel, A. K., & Fries, P. (2010). Beta-band oscillations — signalling the status quo? Current Opinion in Neurobiology, 20(2), 156–165. 10.1016/J.CONB.2010.02.015

Favaro, J., Colombo, M. A., Mikulan, E., Sartori, S., Nosadini, M., Pelizza, M. F., Rosanova, M., Sarasso, S., Massimini, M., & Toldo, I. (2023). The maturation of aperiodic EEG activity across development reveals a progressive differentiation of wakefulness from sleep. NeuroImage, 277, 120264. 10.1016/J.NEUROIMAGE.2023.120264

Ferrarelli, F., Sarasso, S., Guller, Y., Riedner, B. A., Peterson, M. J., Bellesi, M., Massimini, M., Postle, B. R., & Tononi, G. (2012). Reduced Natural Oscillatory Frequency of Frontal Thalamocortical Circuits in Schizophrenia. Archives of General Psychiatry, 69(8), 766–774. 10.1001/ARCHGENPSYCHIATRY.2012.147

Feys, O., & De Tiège, X. (2024). From cryogenic to on-scalp magnetoencephalography for the evaluation of paediatric epilepsy. Developmental Medicine & Child Neurology, 66(3), 298–306. 10.1111/DMCN.15689

Fonov, V. S., Evans, A. C., McKinstry, R. C., Almli, C. R., & Collins, D. L. (2009). Unbiased nonlinear average age-appropriate brain templates from birth to adulthood. NeuroImage, 47, S102. 10.1016/s1053-8119(09)70884-5

Gao, R., Peterson, E. J., & Voytek, B. (2017). Inferring synaptic excitation/inhibition balance from field potentials. NeuroImage, 158, 70–78. 10.1016/J.NEUROIMAGE.2017.06.078

Gogtay, N., Giedd, J. N., Lusk, L., Hayashi, K. M., Greenstein, D., Vaituzis, A. C., Nugent, T. F., Herman, D. H., Clasen, L. S., Toga, A. W., Rapoport, J. L., & Thompson, P. M. (2004). Dynamic mapping of human cortical development during childhood through early adulthood. Proceedings of the National Academy of Sciences of the United States of America, 101(21), 8174–8179. 10.1073/PNAS.0402680101/SUPPL_FILE/02680MOVIE4.MPG

Gómez, C. M., Rodríguez-Martínez, E. I., Fernández, A., Maestú, F., Poza, J., & Gómez, C. (2017). Absolute Power Spectral Density Changes in the Magnetoencephalographic Activity During the Transition from Childhood to Adulthood. Brain Topography, 30(1), 87–97. 10.1007/S10548-016-0532-0/TABLES/3

Grayson, D. S., & Fair, D. A. (2017). Development of large-scale functional networks from birth to adulthood: A guide to the neuroimaging literature. NeuroImage, 160, 15–31. 10.1016/j.neuroimage.2017.01.079

Gross, J., Baillet, S., Barnes, G. R., Henson, R. N., Hillebrand, A., Jensen, O., Jerbi, K., Litvak, V., Maess, B., Oostenveld, R., Parkkonen, L., Taylor, J. R., van Wassenhove, V., Wibral, M., & Schoffelen, J. M. (2013). Good practice for conducting and reporting MEG research. NeuroImage, 65, 349–363. 10.1016/J.NEUROIMAGE.2012.10.001

Hämäläinen, M. S., & Lundqvist, D. (2019). MEG as an enabling tool in neuroscience: Transcending boundaries with new analysis methods and devices. Magnetoencephalography: From Signals to Dynamic Cortical Networks: Second Edition, 3–39. 10.1007/978-3-030-00087-5_81/TABLES/3

Hari, R., & Salmelin, R. (2012). Magnetoencephalography: From SQUIDs to neuroscience: Neuroimage 20th Anniversary Special Edition. NeuroImage, 61(2), 386–396. 10.1016/J.NEUROIMAGE.2011.11.074

He, B. J. (2014). Scale-free brain activity: Past, present, and future. Trends in Cognitive Sciences, 18(9), 480–487. 10.1016/j.tics.2014.04.003

He, W., Brock, J., & Johnson, B. W. (2015). Face processing in the brains of pre-school aged children measured with MEG. NeuroImage, 106, 317–327. 10.1016/J.NEUROIMAGE.2014.11.029

Heinrichs-Graham, E., McDermott, T. J., Mills, M. S., Wiesman, A. I., Wang, Y. P., Stephen, J. M., Calhoun, V. D., & Wilson, T. W. (2018). The lifespan trajectory of neural oscillatory activity in the motor system. Developmental Cognitive Neuroscience, 30, 159–168. 10.1016/J.DCN.2018.02.013

Hill, A. T., Clark, G. M., Bigelow, F. J., Lum, J. A., & Enticott, P. G. (2022). Periodic and aperiodic neural activity displays age-dependent changes across early-to-middle childhood. Developmental Cognitive Neuroscience, 54, 101076. 10.1016/j.dcn.2022.101076

Hill, R. M., Boto, E., Holmes, N., Hartley, C., Seedat, Z. A., Leggett, J., Roberts, G., Shah, V., Tierney, T. M., Woolrich, M. W., Stagg, C. J., Barnes, G. R., Bowtell, R. R., Slater, R., & Brookes, M. J. (2019). A tool for functional brain imaging with lifespan compliance. Nature Communications 2019 10:1, 10(1), 1–11. 10.1038/s41467-019-12486-x

Hill, R. M., Boto, E., Rea, M., Holmes, N., Leggett, J., Coles, L. A., Papastavrou, M., Everton, S. K., Hunt, B. A. E., Sims, D., Osborne, J., Shah, V., Bowtell, R., & Brookes, M. J. (2020). Multi-channel whole-head OPM-MEG: Helmet design and a comparison with a conventional system. NeuroImage, 219, 116995. 10.1016/J.NEUROIMAGE.2020.116995

Hill, R. M., Devasagayam, J., Holmes, N., Boto, E., Shah, V., Osborne, J., Safar, K., Worcester, F., Mariani, C., Dawson, E., Woolger, D., Bowtell, R., Taylor, M. J., & Brookes, M. J. (2022). Using OPM-MEG in contrasting magnetic environments. NeuroImage, 253, 119084. 10.1016/J.NEUROIMAGE.2022.119084

Hoekema, R., Wieneke, G. H., Leijten, F. S. S., Van Veelen, C. W. M., Van Rijen, P. C., Huiskamp, G. J. M., Ansems, J., & Van Huffelen, A. C. (2003). Measurement of the conductivity of skull, temporarily removed during epilepsy surgery. Brain Topography, 16(1), 29–38. 10.1023/A:1025606415858/METRICS

Holmes, N., Tierney, T. M., Leggett, J., Boto, E., Mellor, S., Roberts, G., Hill, R. M., Shah, V., Barnes, G. R., Brookes, M. J., & Bowtell, R. (2019). Balanced, bi-planar magnetic field and field gradient coils for field compensation in wearable magnetoencephalography. Scientific Reports 2019 *9*:1, 9(1), 1–15. 10.1038/s41598-019-50697-w

Hunt, B. A. E., Wong, S. M., Vandewouw, M. M., Brookes, M. J., Dunkley, B. T., & Taylor, M. J. (2019). Spatial and spectral trajectories in typical neurodevelopment from childhood to middle age. Network Neuroscience, 3(2), 497–520. 10.1162/netn_a_00077

Huttenlocher, P. R., & Dabholkar, A. S. (1997). Regional Differences in Synaptogenesis in Human Cerebral Cortex. J. Comp. Neurol, 387, 167–178. 10.1002/(SICI)1096-9861(19971020)387:2

Larsen, B., Cui, Z., Adebimpe, A., Pines, A., Alexander-Bloch, A., Bertolero, M., Calkins, M. E., Gur, R. E., Gur, R. C., Mahadevan, A. S., Moore, T. M., Roalf, D. R., Seidlitz, J., Sydnor, V. J., Wolf, D. H., & Satterthwaite, T. D. (2022). A developmental reduction of the excitation:inhibition ratio in association cortex during adolescence. Science Advances, 8(5), 8750. 10.1126/SCIADV.ABJ8750

Lendner, J. D., Helfrich, R. F., Mander, B. A., Romundstad, L., Lin, J. J., Walker, M. P., Larsson, P. G., & Knight, R. T. (2020). An electrophysiological marker of arousal level in humans. ELife, 9, 1–29. 10.7554/ELIFE.55092

Mahjoory, K., Schoffelen, J. M., Keitel, A., & Gross, J. (2020). The frequency gradient of human resting-state brain oscillations follows cortical hierarchies. ELife, 9, 1–18. 10.7554/ELIFE.53715

Manning, J. R., Jacobs, J., Fried, I., & Kahana, M. J. (2009). Broadband shifts in local field potential power spectra are correlated with single-neuron spiking in humans. Journal of Neuroscience, 29(43), 13613–13620.

Manyukhina, V. O., Prokofyev, A. O., Galuta, I. A., Goiaeva, D. E., Obukhova, T. S., Schneiderman, J. F., Altukhov, D. I., Stroganova, T. A., & Orekhova, E. V. (2022). Globally elevated excitation–inhibition ratio in children with autism spectrum disorder and below-average intelligence. Molecular Autism, 13(1), 1–14. 10.1186/S13229-022-00498-2/TABLES/2

Marshall, P. J., Bar-Haim, Y., & Fox, N. A. (2002). Development of the EEG from 5 months to 4 years of age. Clinical Neurophysiology, 113(8), 1199–1208. 10.1016/S1388-2457(02)00163-3

McIntosh, A. R., Kovacevic, N., Lippe, S., Garrett, D., Grady, C., & Jirsa, V. (2010). The Development of a Noisy Brain. Archives Italiennes de Biologie, 148(3), 323–337. 10.4449/AIB.V148I3.1225

McSweeney, M., Morales, S., Valadez, E. A., Buzzell, G. A., & Fox, N. A. (2021). Longitudinal age- and sex-related change in background aperiodic activity during early adolescence. Developmental Cognitive Neuroscience, 52, 101035. 10.1016/J.DCN.2021.101035

McSweeney, M., Morales, S., Valadez, E. A., Buzzell, G. A., Yoder, L., Fifer, W. P., Pini, N., Shuffrey, L. C., Elliott, A. J., Isler, J. R., & Fox, N. A. (2023). Age-related trends in aperiodic EEG activity and alpha oscillations during early-to middle-childhood. NeuroImage, 269, 119925. 10.1016/J.NEUROIMAGE.2023.119925

Medel, V., Irani, M., Crossley, N., Ossandón, T., & Boncompte, G. (2023). Complexity and 1/f slope jointly reflect brain states. Scientific Reports 2023 *13*:1, *13*(1), 1–12. 10.1038/s41598-023-47316-0

Merkin, A., Sghirripa, S., Graetz, L., Smith, A. E., Hordacre, B., Harris, R., Pitcher, J., Semmler, J., Rogasch, N. C., & Goldsworthy, M. (2023). Do age-related differences in aperiodic neural activity explain differences in resting EEG alpha? Neurobiology of Aging, 121, 78–87. 10.1016/J.NEUROBIOLAGING.2022.09.003

Miller, K. J., Honey, C. J., Hermes, D., Rao, R. P. N., denNijs, M., & Ojemann, J. G. (2014). Broadband changes in the cortical surface potential track activation of functionally diverse neuronal populations. NeuroImage, 85, 711–720. 10.1016/J.NEUROIMAGE.2013.08.070

Miller, K. J., Sorensen, L. B., Ojemann, J. G., & Den Nijs, M. (2009). Power-law scaling in the brain surface electric potential. PLoS Computational Biology, 5(12), e1000609.

Miskovic, V., Ma, X., Chou, C. A., Fan, M., Owens, M., Sayama, H., & Gibb, B. E. (2015). Developmental changes in spontaneous electrocortical activity and network organization from early to late childhood. NeuroImage, 118, 237–247. 10.1016/J.NEUROIMAGE.2015.06.013

Nolte, G. (2003). The magnetic lead field theorem in the quasi-static approximation and its use for magnetoenchephalography forward calculation in realistic volume conductors. Physics in Medicine and Biology, 48(22), 3637–3652. 10.1088/0031-9155/48/22/002

Norbom, L. B., Ferschmann, L., Parker, N., Agartz, I., Andreassen, O. A., Paus, T., Westlye, L. T., & Tamnes, C. K. (2021). New insights into the dynamic development of the cerebral cortex in childhood and adolescence: Integrating macro- and microstructural MRI findings. Progress in Neurobiology, 204, 102109. 10.1016/J.PNEUROBIO.2021.102109

Okada, Y. C., Lahteenmäki, A., & Xu, C. (1999). Experimental analysis of distortion of magnetoencephalography signals by the skull. Clinical Neurophysiology, 110(2), 230–238. 10.1016/S0013-4694(98)00099-6

Olson, I. R., McCoy, D., Klobusicky, E., & Ross, L. A. (2013). Social cognition and the anterior temporal lobes: A review and theoretical framework. Social Cognitive and Affective Neuroscience, 8(2), 123–133. 10.1093/scan/nss119

Oostenveld, R., Fries, P., Maris, E., & Schoffelen, J. M. (2011). FieldTrip: Open source software for advanced analysis of MEG, EEG, and invasive electrophysiological data. Computational Intelligence and Neuroscience, 2011, 1. 10.1155/2011/156869

Ostlund, B., Donoghue, T., Anaya, B., Gunther, K. E., Karalunas, S. L., Voytek, B., & Pérez-Edgar, K. E. (2022). Spectral parameterization for studying neurodevelopment: How and why. Developmental Cognitive Neuroscience, 54, 101073. 10.1016/J.DCN.2022.101073

Pani, S. M., Saba, L., & Fraschini, M. (2022). Clinical applications of EEG power spectra aperiodic component analysis: A mini-review. Clinical Neurophysiology, 143, 1–13. 10.1016/J.CLINPH.2022.08.010

Paolicelli, R. C., Bolasco, G., Pagani, F., Maggi, L., Scianni, M., Panzanelli, P., Giustetto, M., Ferreira, T. A., Guiducci, E., Dumas, L., Ragozzino, D., & Gross, C. T. (2011). Synaptic pruning by microglia is necessary for normal brain development. Science, 333(6048), 1456–1458. 10.1126/SCIENCE.1202529/SUPPL_FILE/PAOLICELLI.SOM.PDF

Parellada, M., Andreu-Bernabeu, Á., Burdeus, M., San José Cáceres, A., Urbiola, E., Carpenter, L. L., Kraguljac, N. V., McDonald, W. M., Nemeroff, C. B., Rodriguez, C. I., Widge, A. S., State, M. W., & Sanders, S. J. (2023). In search of biomarkers to guide interventions in autism spectrum disorder: a systematic review. The American Journal of Psychiatry, 180(1), 23. 10.1176/APPI.AJP.21100992

Partanen, E., Leminen, A., de Paoli, S., Bundgaard, A., Kingo, O. S., Krøjgaard, P., & Shtyrov, Y. (2017). Flexible, rapid and automatic neocortical word form acquisition mechanism in children as revealed by neuromagnetic brain response dynamics. NeuroImage, 155, 450–459. 10.1016/J.NEUROIMAGE.2017.03.066

Pedersen, M., Abbott, D. F., & Jackson, G. D. (2022). Wearable OPM-MEG: A changing landscape for epilepsy. Epilepsia, 63(11), 2745–2753. 10.1111/EPI.17368

Perica, M. I., Calabro, F. J., Larsen, B., Foran, W., Yushmanov, V. E., Hetherington, H., Tervo-Clemmens, B., Moon, C. H., & Luna, B. (2022). Development of frontal GABA and glutamate supports excitation/inhibition balance from adolescence into adulthood. Progress in Neurobiology, 219, 102370. 10.1016/J.PNEUROBIO.2022.102370

Pines, A. R., Larsen, B., Cui, Z., Sydnor, V. J., Bertolero, M. A., Adebimpe, A., Alexander-Bloch, A. F., Davatzikos, C., Fair, D. A., Gur, R. C., Gur, R. E., Li, H., Milham, M. P., Moore, T. M., Murtha, K., Parkes, L., Thompson-Schill, S. L., Shanmugan, S., Shinohara, R. T., … Satterthwaite, T. D. (2022). Dissociable multi-scale patterns of development in personalized brain networks. Nature Communications 2022 *13*:1, *13*(1), 1–15. 10.1038/s41467-022-30244-4

Pozzorini, C., Naud, R., Mensi, S., & Gerstner, W. (2013). Temporal whitening by power-law adaptation in neocortical neurons. Nature Neuroscience 2013 *16*:7, *16*(7), 942–948. 10.1038/nn.3431

Rea, M., Holmes, N., Hill, R. M., Boto, E., Leggett, J., Edwards, L. J., Woolger, D., Dawson, E., Shah, V., Osborne, J., Bowtell, R., & Brookes, M. J. (2021). Precision magnetic field modelling and control for wearable magnetoencephalography. NeuroImage, 241, 118401. 10.1016/J.NEUROIMAGE.2021.118401

Rhodes, N., Rea, M., Boto, E., Rier, L., Shah, V., Hill, R. M., Osborne, J., Doyle, C., Holmes, N., Coleman, S. C., Mullinger, K., Bowtell, R., & Brookes, M. J. (2023). Measurement of Frontal Midline Theta Oscillations using OPM-MEG. NeuroImage, 271, 120024. 10.1016/J.NEUROIMAGE.2023.120024

Richards, J. E., Sanchez, C., Phillips-Meek, M., & Xie, W. (2016). A database of age-appropriate average MRI templates. NeuroImage, 124, 1254–1259. 10.1016/J.NEUROIMAGE.2015.04.055

Rier, L., Rhodes, N., Pakenham, D. O., Boto, E., Holmes, N., Hill, R. M., Reina Rivero, G., Shah, V., Doyle, C., Osborne, J., Bowtell, R. W., Taylor, M., & Brookes, M. J. (2024). Tracking the neurodevelopmental trajectory of beta band oscillations with optically pumped magnetometer-based magnetoencephalography. ELife, 13. 10.7554/ELIFE.94561

Roberts, G., Holmes, N., Alexander, N., Boto, E., Leggett, J., Hill, R. M., Shah, V., Rea, M., Vaughan, R., Maguire, E. A., Kessler, K., Beebe, S., Fromhold, M., Barnes, G. R., Bowtell, R., & Brookes, M. J. (2019). Towards OPM-MEG in a virtual reality environment. NeuroImage, 199, 408–417. 10.1016/J.NEUROIMAGE.2019.06.010

Rodríguez-Martínez, E. I., Ruiz-Martínez, F. J., Barriga Paulino, C. I., & Gómez, C. M. (2017). Frequency shift in topography of spontaneous brain rhythms from childhood to adulthood. Cognitive Neurodynamics, 11(1), 23–33. 10.1007/S11571-016-9402-4/FIGURES/6

Rosanova, M., Casali, A., Bellina, V., Resta, F., Mariotti, M., & Massimini, M. (2009). Natural Frequencies of Human Corticothalamic Circuits. Journal of Neuroscience, 29(24), 7679–7685. 10.1523/JNEUROSCI.0445-09.2009

Safar, K., Vandewouw, M., Sato, J., Devasagayam, J., Hill, R., Rea, M., Brookes, M., & Taylor, M. (2024). Using optically pumped magnetometers to replicate task-related responses in next generation magnetoencephalography. Scientific Reports, 14, 6513. doi.org/10.21203/rs.3.rs-3263385/v1

Schäfer, C. B., Morgan, B. R., Ye, A. X., Taylor, M. J., & Doesburg, S. M. (2014). Oscillations, networks, and their development: MEG connectivity changes with age. Human Brain Mapping, 35(10), 5249–5261. 10.1002/hbm.22547

Schaworonkow, N., & Voytek, B. (2021). Longitudinal changes in aperiodic and periodic activity in electrophysiological recordings in the first seven months of life. Developmental Cognitive Neuroscience, 47, 100895. 10.1016/J.DCN.2020.100895

Seedat, Z. A., Pier, K. St., Holmes, N., Rea, M., Al-Hilaly, L., Tierney, T. M., Embury, C. M., Pardington, R., Mullinger, K. J., Cross, J. H., Boto, E., & Brookes, M. J. (2024). Simultaneous whole-head electrophysiological recordings using EEG and OPM-MEG. Imaging Neuroscience, 2, 1–15. 10.1162/IMAG_A_00179

Segalowitz, S. J., Santesso, D. L., & Jetha, M. K. (2010). Electrophysiological changes during adolescence: A review. Brain and Cognition, 72(1), 86–100. 10.1016/J.BANDC.2009.10.003

Seymour, R. A., Alexander, N., Mellor, S., O’Neill, G. C., Tierney, T. M., Barnes, G. R., & Maguire, E. A. (2021). Using OPMs to measure neural activity in standing, mobile participants. NeuroImage, 244, 118604. 10.1016/J.NEUROIMAGE.2021.118604

Sydnor, V. J., Larsen, B., Bassett, D. S., Alexander-Bloch, A., Fair, D. A., Liston, C., Mackey, A. P., Milham, M. P., Pines, A., Roalf, D. R., Seidlitz, J., Xu, T., Raznahan, A., & Satterthwaite, T. D. (2021). Neurodevelopment of the association cortices: Patterns, mechanisms, and implications for psychopathology. Neuron, 109(18), 2820–2846. 10.1016/J.NEURON.2021.06.016

The Mathworks Inc. (2018). MATLAB. In www.mathworks.com/products/matlab (2021a).

Thuwal, K., Banerjee, A., & Roy, D. (2021). Aperiodic and Periodic Components of Ongoing Oscillatory Brain Dynamics Link Distinct Functional Aspects of Cognition across Adult Lifespan. ENeuro, 8(5). 10.1523/ENEURO.0224-21.2021

Tierney, T. M., Alexander, N., Mellor, S., Holmes, N., Seymour, R., O’Neill, G. C., Maguire, E. A., & Barnes, G. R. (2021). Modelling optically pumped magnetometer interference in MEG as a spatially homogeneous magnetic field. NeuroImage, 244, 118484. 10.1016/J.NEUROIMAGE.2021.118484

Tikhonov, A. N. (1943). On the stability of inverse problems. Comptes Rendus De L Academie Des Sciences De L Urss, 39, 176–179.

Tröndle, M., Popov, T., Dziemian, S., & Langer, N. (2022). Decomposing the role of alpha oscillations during brain maturation. ELife, 11. 10.7554/ELIFE.77571

Tzourio-Mazoyer, N., Landeau, B., Papathanassiou, D., Crivello, F., Etard, O., Delcroix, N., Mazoyer, B., & Joliot, M. (2002). Automated anatomical labeling of activations in SPM using a macroscopic anatomical parcellation of the MNI MRI single-subject brain. NeuroImage, 15, 273–289. 10.1006/nimg.2001.0978

Usher, M., Stemmler, M., & Olami, Z. (1995). Dynamic Pattern Formation Leads to Noise in Neural Populations. Physical Review Letters, 74(2), 326. 10.1103/PhysRevLett.74.326

Van Veen, B. B. D., Van Drongelen, W., Yuchtman, M., & Suzuki, A. (1997). Localization of brain electrical activity via linearly constrained minimum variance spatial filtering. IEEE Transactions on Biomedical Engineering, 44(9), 867–880. 10.1109/10.623056

Vanderwal, T., Kelly, C., Eilbott, J., Mayes, L. C., & Castellanos, F. X. (2015). Inscapes: A movie paradigm to improve compliance in functional magnetic resonance imaging. NeuroImage, 122, 222–232. 10.1016/j.neuroimage.2015.07.069

Voytek, B., & Knight, R. T. (2015). Dynamic Network Communication as a Unifying Neural Basis for Cognition, Development, Aging, and Disease. Biological Psychiatry, 77(12), 1089–1097. 10.1016/J.BIOPSYCH.2015.04.016

Voytek, B., Kramer, M. A., Case, J., Lepage, K. Q., Tempesta, Z. R., Knight, R. T., & Gazzaley, A. (2015). Age-Related Changes in 1/f Neural Electrophysiological Noise. Journal of Neuroscience, 35(38), 13257–13265. 10.1523/JNEUROSCI.2332-14.2015

Waschke, L., Donoghue, T., Fiedler, L., Smith, S., Garrett, D. D., Voytek, B., & Obleser, J. (2021). Modality-specific tracking of attention and sensory statistics in the human electrophysiological spectral exponent. ELife, 10. 10.7554/ELIFE.70068

Whitford, T. J., Rennie, C. J., Grieve, S. M., Clark, C. R., Gordon, E., & Williams, L. M. (2007). Brain maturation in adolescence: Concurrent changes in neuroanatomy and neurophysiology. Human Brain Mapping, 28(3), 228–237. 10.1002/HBM.20273

Wittevrongel, B., Holmes, N., Boto, E., Hill, R., Rea, M., Libert, A., Khachatryan, E., Van Hulle, M. M., Bowtell, R., & Brookes, M. J. (2021). Practical real-time MEG-based neural interfacing with optically pumped magnetometers. BMC Biology, 19(1), 1–15. 10.1186/S12915-021-01073-6/FIGURES/4

Zetter, R., Iivanainen, J., & Parkkonen, L. (2019). Optical Co-registration of MRI and On-scalp MEG. Scientific Reports 2019 *9*:1, *9*(1), 1–9. 10.1038/s41598-019-41763-4

